# Invasion but not hybridisation is associated with ecological niche shift in monkeyflowers

**DOI:** 10.1101/688515

**Authors:** Daniele Da Re, Angel P. Olivares, William Smith, Mario Vallejo-Marín

## Abstract

**Background:** The ecological niche occupied by novel hybrids can influence their establishment as well as the potential to coexist with their parents. Hybridisation generates new phenotypic combinations, which, in some cases, may allow them to occupy ecological niches outside the environmental envelope of parental taxa. In other cases, hybrids may retain similar ecological niches to their parents, resulting in competition and affecting their coexistence. To date, few studies have quantitatively assessed niche shifts associated with hybridisation in recently introduced populations while simultaneously characterising the niche of parental species in both native and introduced ranges.

**Aims:** In this study, we compared the ecological niche of a novel hybrid plant with the niches of its two parental taxa in the non-native geographic range. We also characterised and compared the parental taxa’s ecological niche of native and introduced populations in order to assess potential niche changes during the invasion process independent of hybridisation.

**Methods:** We studied monkeyflowers (*Mimulus spp.*, Phrymaceae) that were introduced from the Americas to Europe and New Zealand in the last 200 years. We focused on a novel hybrid, triploid, asexual taxon (*M. × robertsii*) that occurs only in the British Isles where its two parents (*M. guttatus* and *M. luteus*) come into secondary contact. We assembled more than 12,000 geo-referenced occurrence records and eight environmental variables of the three taxa across native and introduced ranges, and conducted ecological niche model analysis using maximum entropy, principal component and niche dynamics analysis.

**Results:** We found no evidence of niche shift in the hybrid, *M. × robertsii* compared to introduced populations of both of their parental taxa. The hybrid had a niche more similar to *M. luteus*, which is also the rarest of the parental taxa on the introduced range. Among parental monkeyflowers, *M. guttatus* showed niche conservatism in introduced populations in Europe, but a niche shift in New Zealand, while *M. luteus* showed a niche shift in Europe. However, the evidence of niche shift should be treated with caution due to the occurence of non-analog climatic conditions, small population size and unfilling niche dynamics.

**Conclusions:** Our results suggest that hybridisation in non-native monkyeflowers did not result in a shift in ecological niche. This niche conservation could create competition between parental and derived taxa, the outcome of which will depend on relative competitive abilities. Further work is needed to establish if the expansion of the hybrid in the introduced range is causally related to the apparent rarity of one of the parents (*M.luteus*). Finally, the comparison of native and non-native populations of parental taxa, suggest that whether invasions result in niche shifts or not depends on both taxon and geographic region, highlighting the idiosyncratic nature of biological invasions.

## Introduction

During the Anthropocene, human trade and travel has dispersed species beyond their native range, sometimes connecting previously isolated taxa. Some non-native species represent a threat to biodiversity, human health and economy (Mack et al. 2000; Simberloff et al. 2013, Pysek et al 2017). Understanding non-native species ecology and the potential differences between populations in native and invaded ranges can help us deal with biological invasion and develop effective management strategies. A powerful tool to characterise the broad scale environmental conditions in which native and non-native populations occur is niche modelling (Guisan et al. 2017). Ecological niche models (ENMs; Anderson 2012) are correlative statistical techniques that estimate the relationship between geo-referenced occurrences and environmental variables, allowing then to predict the suitable habitat distribution at large geographic scale (Peterson et al. 2003). ENMs are widely used to predict invaded areas projecting models fitted on the empirical native species’ distribution (Guisan et al. 2017). ENMs can also be used to quantify niche changes between different areas (e.g., native and introduced), by comapring niche differences in the environmental space rather than in the projected geographical one (Warren et al. 2008; Broenniman et al. 2012). Assuming that a species colonized all the suitable environment in the native range, Petitpierre et al. (2012) described two processes that could differentiate native and non-native populations niches: (1) Niche expansion (i.e., species colonizing novel environmental condition in the invaded area resulting from adaptation to novel local conditions) and (2) niche unfilling (i.e. a partial filling of the native niche in the invaded range). Assessing whether these processes lead to significant (realized) niche differentiation between native and non-native populations entails testing two different hypotheses, namely niche equivalency (native and non-native niches are indistinguishable and interchangeable) and niche similarity (whether niches are more similar than expected by chance; Warren et al. 2008). Comparisons between introduced and native populations enables testing the extent to which local adaptation (niche expansion) or niche matching (niche unfilling) helps explaining the realized niche of non-native populations.

In addition to the potential occupation of new ecological spaces, another consequence of biological invasions is the hybridization of species as previously isolated taxa come into secondary contact. Hybridization can form organisms that are genetically more diverse than their parental taxa and in some cases result in novel taxa (Dietz and Edwards 2006; Marchant et al. 2016; Parisod and Broennimann 2016; Vallejo-Marín and Hiscock 2016; Visger et al. 2016; Molina-Henao and Hopkins 2019). The new genotypes and phenotypes created through hybridization can potentially enable hybrid taxa to exploit new environmental conditions compared to their parental taxa (Sheth and Anger 2014), thus potentially shifting their fundamental niche (Marchant et al. 2016; Parisod and Broennimann 2016). However, to date only few studies have investigated the extent to which hybridisation vs. range expansion is associated with shifts in niche occupancy (e.g. Mukherjee et al. 2012; Thornton and Murray 2014, Visger et al. 2016; Molina-Henao and Hopkins 2019).

Some species of monkeyflowers (*Mimulus spp.*) are a prime example of recent plant invasion and hybridization events that have yielded widespread, novel hybrids that exist only in the non-native range (Stace 2010; Stace et al. 2015). Among these hybrid taxa, probably the best-studied case is the triploid hybrid *M. × robertsii* in the United Kingdom (UK). The hybrid monkeyflower, *M. × robertsii* is the product of crosses between two non-native species that are completely allopatric in their native range: the tetraploid *M. luteus* from South America (Chile and Argentina), and the diploid *M. guttatus* from western North America (Mexico to Alaska). Both of the parental species were introduced in Europe in the 19^th^ century, and were used in the horticultural trade probably due to their striking yellow and red flowers. In the UK, *M. guttatus* was introduced in 1812, after which it became naturalised and is currently widely distributed throughout Great Britain and northern Ireland. *M. guttatus* has also been introduced into mainland Europe, Iceland, the Faroe Islands, New Zealand and eastern North America. The introduction of *M. guttatus* to New Zealand appears to date back at least to 1878 (Owen 1996), while the introduction history in other regions is less well known. The South American *M. luteus* appears to have arrived to the UK a few decades later with records dating back to at least the 1830s. Historical records suggest that *M. luteus* has been found across the UK and in other areas of Europe and New Zealand. At present, naturalised populations of *M. luteus* are very rare compared to other non-native monkeyflowers and are mainly restricted to the UK (Vallejo-Marín and Lye 2013). The origin and exact parentage of *M. × robertsii* is unknown, but naturalised populations of these hybrids became established by 1844 and since then, this taxon has become widely distributed in the UK (Stace et al. 2015), with about 40% monkeyflower populations being composed partially or entirely of hybrids (Vallejo-Marin and Lye 2013). Both hybrid and parental taxa occupy mainly wet habitats such as banks of streams and rivers, bogs and other wet places (Truscott et al. 2006). To date, no study has been conducted characterising the ecological niche of non-native and hybrid populations of monkeyflowers.

In this study we compare ecological niches between parental and hybrid monkeyflowers and among native and non-native populations of the parental taxa. Specifically, we address the following questions: 1) Does the ecological niche of parental taxa shift during the invasion process, and if so to what extent? 2) Which regions in the native range have the highest ecological niche similarity to the conditions faced by introduced populations in invaded areas? 3) Does the hybrid fundamental niche differs from the parental native and invaded fundamental niche?

## Materials and Methods

### Geo-referenced occurrences

Geo-referenced occurrences data of the three taxa and their subordinates taxonomic ranks were downloaded from the Global Biodiversity Information Facility (GBIF 2016; www.gbif.org), the Nodo Nacional de Información de Biodiversidad (GBIF España 2016; www.gbif.es), the GBIF France (GBIF France 2016; www.gbif.fr), the Botanical Society of Britain and Ireland (BSBI 2016; www.bsbi.org), the NBN gateway (NBN 2016; https://data.nbn.org.uk), the FloraWeb (FloraWeb 2016; www.floraweb.de), the Integrated Digitized Biocollections (iDigBio 2016; www.idigbio.org) and the Kasviatlas (Lampien and Lahti 2016; http://www.luomus.fi/kasviatlas). In addition to these sources, records of *M. guttatus* from its native range were included from Oneil (2014).

Records with erroneous coordinates (e.g., records located in sea), expressed with different geographic coordinates than latitude and longitude decimal degrees and with a coordinate accuracy higher than 1km were excluded. In order to make sure that the species occurrences were encompassed in the time span of the environmental variables, only data from the year 1950 onwards was kept.

### Environmental variables

30 arc sec spatial resolution bioclimatic variables describing the current environmental conditions (1950 - 1990 year span) were downloaded from the WorldClim database (Hijman et al. 2005; www.worldclim.org) and managed using R v3.4.0 (R Core Team 2019). In accordance with previous studies on native populations of monkeyflowers (Grossenbacher et al. 2014; Sobel 2014), eight of the most important bioclimatic variables for the niche of *Mimulus ssp.* were chosen for analysis. These bioclimatic variables were cropped to the distribution of the outermost records of each taxon plus a buffer of 2º (Table 1; cf. Sobel 2014).

**Table 1.** List of variables used to model.

| Variables | Abbreviation |
| --- | --- |
| Annual mean temperature | Bio1 |
| Temperature seasonality | Bio4 |
| Maximum temperature of warmest month | Bio5 |
| Minimum temperature of coldest month | Bio6 |
| Annual mean precipitation | Bio12 |
| Precipitation seasonality | Bio15 |
| Precipitation of wettest quarter | Bio16 |
| Precipitation of driest quarter | Bio17 |

### Niche analysis

Since niche differentiation in environmental space may or may not translate into occupation of different geographic space (Warren et al. 2008), all the analysis were computed in the environmental space among the three species in both native and invasive range using *ecospat* R package. The ecological niche space occupied by each species in each native/invaded area was investigated using environmental PCA (PCA-env, Broennimann et al. 2012). PCA-env is an ordination technique calibrated on the whole environmental space of both the native and the invaded study areas, which allows to plot a kernel smoothed density of occurrences for each species in the principal component space (Cola et al. 2017). In order to prevent projecting a model in non-analog climatic conditions (a combination of climatic conditions that are not found in the climatic envelope of the areas/time where the model is trained), we computed a PCA of the environmental predictors between each ranges and check their overlap (Guisan et al. 2017).

The overlap between two different niches in the ecological space was quantified using Schooner’s D metric (Warren et al. 2008), which ranges from no overlap (D = 0) to complete overlap (D = 1). Additionally, the niche overlap can be decomposed into niche unfilling and niche expansion. While the former reports the proportion of native species occurrences densities that have environmental conditions different from those in the non-native-range, the latter asses the contrary, indicating the proportion of non-native occurrences having environmental conditions different from the native one. This decomposition provides more information about the drivers of niche dynamic between native and invaded ranges (Petitpierre et al. 2012; Guisan et al. 2014), or about how two sister species have evolved different niches. Each indexes was computed using the 90th percentile of the available environmental conditions common to both ranges, in order to remove the marginal environments and avoids bias due to artefacts of the density function (Petitpierre et al. 2012; Cola et al. 2017; Villaverde et al. 2017).

In addition, we computed niche equivalency and niche similarity tests (Warren et al. 2008) to assess the statistical significance difference between estimated realized niches. We tested niche divergence (alternative = “lower”) for both analysis and we randomly shifted the invasive niche only in the comparisons between native and invasive niche (rand.type = 2). Niche equivalence tests assess whether the realized ecological niches of two taxa are environmentally identical and interchangeable. For each taxa, it test whether the observed D derived from taxa’s occurrences is constant when the occurrences of both taxa are randomly reallocated and compared to a null distribution generated by 100 pseudoreplicate datasets (Warren et al. 2008; Broennimann et al. 2012). The hypothesis of niche equivalency is rejected when observed values of D are significantly different (p < 0.05) from the simulated values and so the taxa do not have equivalent realized niches. The niche equivalency test is often rejected because it uses species’ occurrences only and does not account for occurrences’ surrounding space. For these reasons, other authors (e. g. Hu et al. 2016) suggested to use this test for evaluating the transferability of niche models in space and time only and to assess biogeographic hypothesis using the niche similarity test (Peterson 2011). In fact, the niche similarity test assesses if the ecological niches of two taxa are more similar than expected by chance, accounting for the differences in the surrounding environmental conditions in the geographic areas where both species are distributed (Warren et al. 2010). It evaluates whether the overlap between observed niches in two ranges is different from the overlap between the observed niche in one range and niches selected at random from the other range (Warren et al. 2008; Broennimann et al. 2012). The niche similarity test indicates niche similarity while accounting for the similarity in background environmental conditions.

### Ecological Niche Modelling (ENM)

Ecological niche models were performed using Maxent v3.4 (Phillips et al. 2017) through the R package *dismo* (Hijmans et al. 2017). To reduce the effects of sampling bias and thus avoid a possible source of model inaccuracy (Phillips et al. 2006; Phillips et al. 2009; Syfert et al. 2013), spatial filtering with a thinning distance of 2 km was applied to the final dataset of the three species using the R package *spThin* (Aiello-Lammens et al. 2015), while in order to avoid overfitting, species-specific tuning of Maxent models’ settings using AICc was investigated using the R package *ENMeval* (Muscarella et al. 2014). The performed models were built and evaluated at the occurrences geographical space plus a buffer of 2º for each species (Sobel 2014; Soberon 2018), and then were re-projected into the environmental conditions of their respective native/invasive population or vice versa. Nevertheless, to restrict the modelling to the conditions encountered in the original range, extrapolation was not applied and clamping was done when projecting. Models were set up to obtain a logistic response of the predicted distribution and were evaluated using the area under the curve (AUC) provided for the test data (Phillips, Anderson and Schapire 2006; Ward 2007). AUC values range from 0 to 1. According to the classification of Swets (1988), model with AUC = 0.5 do not discriminate between suitable and unsuitable cells better than a random model, an AUC score >0.7 shows a ‘‘useful’’ discrimination ability, >0.8 shows a ‘‘good’’ model performance and >0.9 ‘‘very good’’ model performance. Recently many authors (Breiner et al. 2015, Cola et al. 2017) suggested to use the Boyce index, a presence-only and threshold-independent evaluator for SDMs ’predictions (Hirzel et al. 2006), for models evaluation in addition to the common used AUC. The Boyce index, computed through ecospat R package (Cola et al. 2017), ranges between – 1 (the model predict areas where presences are more frequent as being highly suitable for the species) and + 1 (the model predictions are consistent with the distribution of presences in the evaluation data set). Values close to zero mean that the model is not different from a random model (Hirzel et al. 2006).

### ENMs projections

The ENMs were trained in each species native and invaded regions and then projected: (1) Native range projected into the invasive range (prospective niche modelling). *M. guttatus*’ western North America occurrences were used to train the native niche model and then projected it into the invasive regions (Europe and New Zealand). Western South America occurrences were used to train *M. luteus* model trained in the native region and then projected into Europe only; (2) Invasive range projected into the native range (retrospective niche modelling). We used the occurrence records from the invasive range (Europe and New Zealand for *M. guttatus*, Europe only for *M. luteus*), and projected it back into western North America and South America, respectively. These analyses show the predicted niche suitability of the native range, based on the estimated ecological niche inferred from a given invasive region; (3) Invasive hybrid niche model projected onto the two parental taxa native range, in order to highlight the predicted niche suitability of the hybrid in the parental taxa native regions.

## Results

A total of 12,480 records were kept after curating the worldwide distributed data. Spatial filtering yielded a final number of 9,079 records across all taxa and geographic regions (Table 2). The number of spatially filtered records per taxon and region varied widely. The taxon with the largest number of records across all regions was *M. guttatus* (6,648) with approximately 73% of those records found in the introduced European range, mostly the Britain and Ireland, and 25% (1,763) in the native North American range. In contrast, after spatial filtering we obtained only 19 records (<1%) in the introduced New Zealand range. There were considerably fewer records of *M. luteus*, with most found in the introduced range (625 or 95% of the total), and only 30 in the native South American range. We had a relatively large number of records of the hybrid *M × robertsii* (1,776) all restricted to Britain and Ireland.

**Table 2.** Reports the models with the lowest AICc selected for each species and their characteristics. All models show good AUC and Boyce index scores.

| Species | Training region | n° of records | Model features | Beta multiplier | AUC<br>(± SD) | Boyce Index |
| --- | --- | --- | --- | --- | --- | --- |
| *M. guttatus* | NA | 1763 | LQP | 1 | $0.819 \pm 0.006$ | 0.999 |
| | EU | 4866 | LQPH | 0.5 | $0.807 \pm 0.002$ | 0.998 |
| | NZ | 19 | LQ | 1 | $0.650 \pm 0.062$ | 0.783 |
| *M. luteus* | SA | 30 | L | 0.5 | $0.867 \pm 0.082$ | 0.924 |
| | EU | 625 | LQPH | 2 | $0.902 \pm 0.012$ | 0.994 |
| *M. robertsii* | EU | 1776 | LQPH | 2 | $0.792 \pm 0.009$ | 0.985 |

Only the models trained in South America and New Zealand used linear and quadratic features exclusively, suggesting that the model complexity increased as the sample size increased (Table 2). Also the AUC metrics were influenced by the sample size, in fact higher scores were obtained by the models having larger sample size (Table 2). Models’ Boyce index values were always values above 0.7, confirming good model performances.

### Principal component analysis and niche similarity

The PCA made on the climatic conditions present in the ranges of *M. guttatus* showed analog climate conditions between the North American and European range (SM1a). On the contrary, non-analog climate and divergent patterns were observed between the North American and New Zealand’s ranges and between Europe and New Zealand’s ones as well (SM1b, c). In the case of *M. luteus*, non-analog climate and divergent patterns were observed between *M. luteus* Nat. range and the European one, thus preventing reprojection (SM2a). However, analog conditions were found between *M. luteus* Nat. range and *M. guttatus* Nat. one (SM2b). According to this findings, only the reprojection of *M. guttatus* Nat. niche into Europe and vice versa was possible.

*M. guttatus* showed relatively low niche overlap between native and non-native ranges (D = 0.190 and D = 0.203, for Europe and New Zealand, respectively). Similarly, niche overlap between the two invaded regions (Europe and New Zealand) was very low (D = 0.043) (Table 3). Low niche overlap is related to niche unfilling in native and introduced regions, while between Europe and New Zealand is associated with niche expansion as indicated by the niche dynamics statistics (Table 3). Evidences of niche conservatism (niches equivalent and more similar than by chance) did not emerge from equivalency and similarity test results between the native niche and the two invasive niches (Table 3). In fact, *M. guttatus* Nat. niche was equivalent but similar by chance to the European populations’ niche and the Nat. niche was not equivalent and similar by chance to the New Zealand one. Regarding the comparison between the two invasive niches, they resulted not equivalent and similar by chance. Low niche overlap (D = 0.309) was observed in the comparison between *M. luteus* Nat. and its european Inv. niche. As evidence of low niche overlap and lack of niche conservatism, both niche unfilling and expansion were observed and the niches equivalency and similarity test resulted in not equivalent and similar by chance niches (Table 3). In the European range, *M. guttatus* Inv. showed high niche similarity (D = 0.734) and niche conservatism with *M. luteus* Inv., having the two niches equivalent and more similar than by chance (Table 3). In contrast, *M. luteus* Nat. niche showed low niche overlap (D = 0.384) and niche expansion when compared to *M. guttatus* Nat. Evidences of niche conservatism came from the comparisons between the parental taxas and the hybrid in their European ranges. European *M. guttatus* Inv. showed high niche similarity (D = 0.606) and non-equivalent but more similar than by chance niches (Table 3). *M. luteus* Inv. showed higher niche overlap with *M. robertsii* (D = 0.705) and niche conservatism, having the two niches equivalent and more similar than by chance (Table 3).

**Table 3.** Results of the niche equivalency and similarity test carried in the environmental space. Signif. codes: P < 0.001 ‘***’; P < 0.01 ‘**’; P < 0.05 ‘*’; ‘ns’ not significant.

| Species pair | Populations | rand.type | Schoener's *D* | Equivalency | Similarity | Unfilling | Expansion | Interpretation |
| --- | --- | --- | --- | --- | --- | --- | --- | --- |
| *M. guttatus* - *M. guttatus* | Nat. (NA) - Inv. (EU) | 2 | 0,190 | ns | ns | **0,612** | 0,000 | Equivalent but similar by chance |
|  | Nat. (NA) - Inv. (NZ) | 2 | 0,203 | ** | ns | **0,616** | 0,006 | Not equivalent and similar by chance, supposed niche divergence |
|  | Inv. (EU) - Inv. (NZ) | 1 | 0,043 | ** | ns | **0,243** | **0,483** | Not equivalent and similar by chance, supposed niche divergence |
| *M. luteus* - *M. luteus* | Nat. (SA) - Inv. (EU) | 2 | 0,309 | * | ns | **0,348** | **0,162** | Not equivalent and similar by chance, supposed niche divergence |
| *M. luteus* - *M. guttatus* | Nat. (SA) - Inv. (EU) | 1 | 0,384 | ns | ** | 0,001 | **0,340** | Equivalent and more similar than by chance, evidences of niche conservatism |
|  | Inv. (EU) - Inv. (EU) | 1 | 0,734 | ns | ** | 0,013 | 0,027 | Equivalent and more similar than by chance, evidences of niche conservatism |
| *M. robertsii* - *M. guttatus* | Nat. (EU) - Inv. (EU) | 1 | 0,606 | ** | ** | 0,049 | 0,000 | Not equivalent but more similar than by chance, there is no niche conservatism but there are similarities |
| *M. robertsii* - *M. luteus* | Nat. (EU) - Inv. (EU) | 1 | 0,705 | ns | ** | 0,055 | 0,000 | Equivalent and more similar than by chance, evidences of niche conservatism |

### Environmental Niche Modelling

*M. guttatus* Nat. ENM trained in North America showed high niche suitability in south-western United States, north western Mexico and the Alaskan coast (Fig. 1a), accordingly to its current distribution. In particular, this model predicted suitable areas close to the Haida Gwaii (Queen Charlotte) Islands and Prince of Wales Island in British Columbia (Canada) and further north and west in Alaska from the southeast of Kodiak Island, and onto the Aleutian Islands range from around Attu Island in the east to Unalaska in the west. Interestingly, the Alaskan coast is also one of few and geographically restricted regions with relatively high niche suitability predicted by the European invasive population’ ENM re-projected into the native range (Fig. 1b). Native population’s ENM re-projected into the invasive European range showed high niche suitability in almost all of the current distribution of *M. guttatus* in western Europe (Fig. 2a). However the predicted suitable area is wider than the one predicted by the European *M. guttatus* Inv.’s ENM, which showed the highest suitability in the British Isles, the north coast of France, parts of Belgium and the Netherlands and the central Germany(Fig. 2b). New Zealand *M. guttatus* (Inv.)’s ENM located suitable areas mainly in the coastal areas and in the northern island (Fig. 2c).

**Figure 1.**
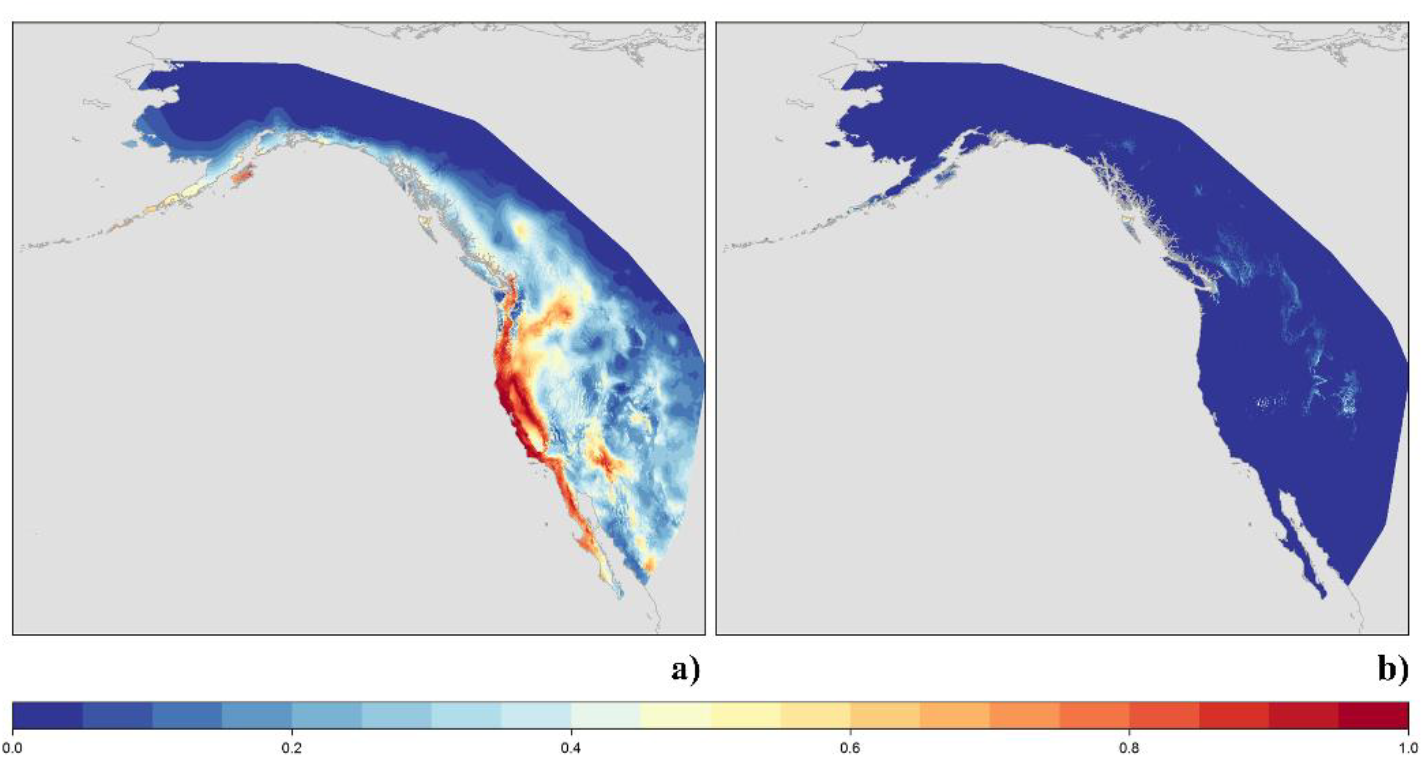
**a, b.** ENM trained on a) the current *M. guttatus* native distribution in North America and b) the current *M. guttatus* invasive distribution in Europe and projected into the native geographical area.

**Figure 2.**
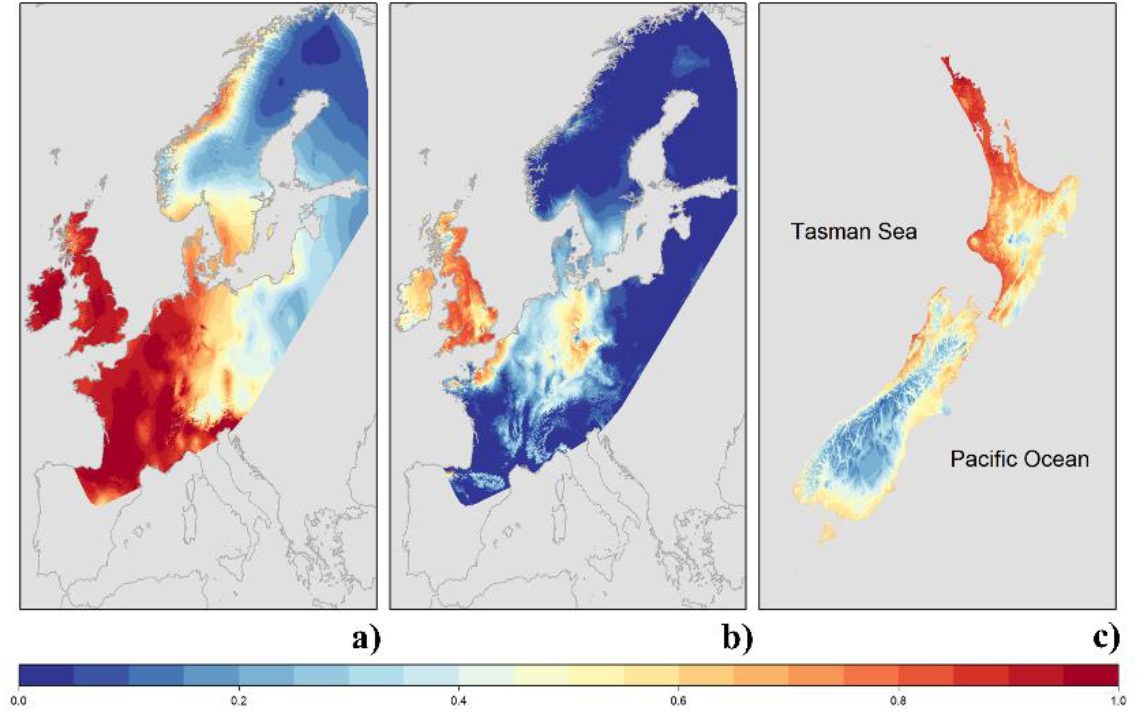
**a, b, c.** ENM trained on a) the current *M. guttatus* native distribution in North America and projected into Europe, b) the current *M. guttatus* invasive distribution in Europe, c) the current *M. guttatus* invasive distribution in New Zealand.

*M. luteus* (Nat.)’ ENM predicted suitable conditions in the southern central Andean region of Chile (Fig. 3a). In Europe, the model trained on invasive population predicted suitable areas mainly in the British Isles, except the south – east England and the Scottish Highlands (Fig. 3b), in accordance with its current distribution. *M. robertsii* ENM showed a highly suitable areas mainly in the British Isles (Fig. 4c) and their predicted distribution resemble the distribution of *M. luteus*’ one (Fig. 4b), contrary to *M. guttatus* that seemed to have a wider distribution even outsite the United Kingdom (UK; Fig. 4c).

**Figure 3.**
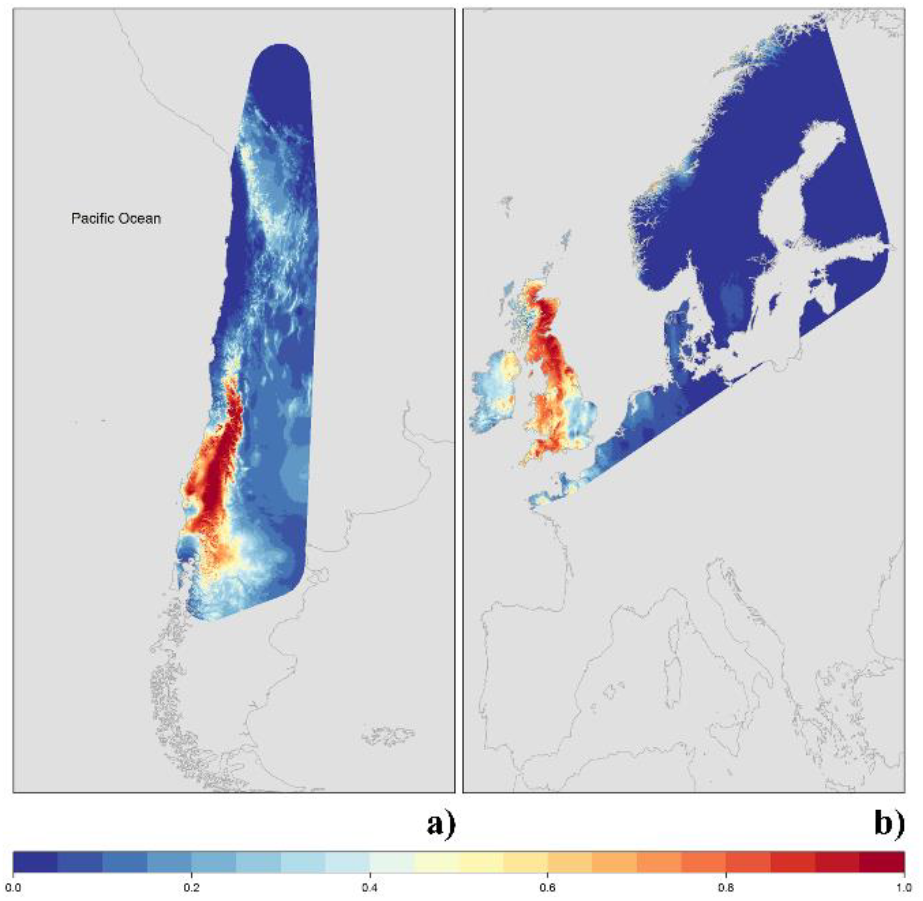
**a, b.** ENM trained on a) the current *M. luteus* native distribution in South America and b) the current *M. guttatus* invasive distribution in Europe.

**Figure 4.**
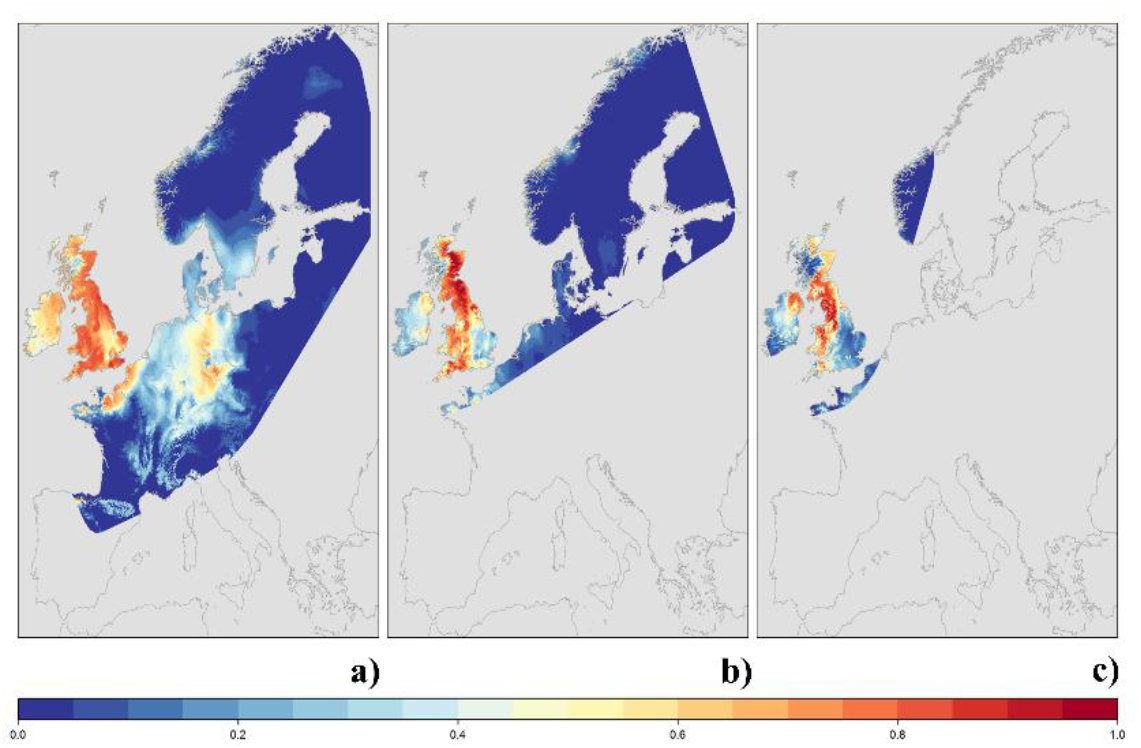
**a, b, c.** ENM trained on the current distribution of the three species in Europe: a) *M. guttatus*, b) *M. luteus*, c) *M. robertsii*.

## Discussion

In this study we modelled and compared the ecological niche of *M. guttatus* and *M. luteus* in their native and invasive ranges, as well as the ecological niche of their hybrid, *M. × robertsii*. While previous studies have analysed the niche of *M. guttatus* using either a correlative (Ferris et al. 2014; Grossenbacher et al. 2014) or a mechanistic approach (Sheth and Angert 2014), our study is the first effort to model the ecological niche and spatial distribution of the Southamerican taxon *M. luteus* and the hybrid *M. × robertsii.* Furthermore, our study allowed us to compare the ecological niches of these three closely related taxa using and measure niche differences in a gridded environmental space built choosing ecologically relevant variables (Early and Sax 2014). Below, we discuss how the niche models produced here can be used to understand potential shifts in ecological niche following hybridisation, as well as the changes in niche when associated with range expansion and biological invasions.

### Ecological niche of the hybrid

One of the main goals of our study was to determine if a novel hybrid occupied a new ecological niche relative to its parents. In general, the ecological niche of the hybrid *M. × robertsii* is similar to both parental taxa, showing a high overlap in the environmental space (Figure 5). However, the the ecological niche of *M. × robertsii* is statistically different from invasive populations of *M. guttatus* indicating that hybridisation contributes to niche differentiation from at least one parental taxon. The comparison of ecological niche between the hybrid and each parental taxon suggests that the niche of *M. × robertsii* is equivalent and more similar to the *M. luteus* than to *M. guttatus*. The asymmetry of niche similarity between the hybrids and each parental taxon may translate in different probabilities of co-occurrence and competition (Costa and Schlupp 2012; Mukherjee et al. 2012; Molina-Henao and Hopkins 2019). The co-occurrence of *M. luteus* and the hybrid may provide more opportunities for competition between these two taxa. If the hybrid is a more aggressive competitor than the South American parent, it is possible that competitive interactions may help to explain the apparent decline in occurrence of *M. luteus* compared to the hybrids. Biotic interactions are important in the successful establishment of hybrids in the same environment of its parental taxa (Gaskin 2016; Marchant et al. 2016), and may also be responsible in shaping the ecological sorting of invasive monkeyflowers.

**Figure 5.**
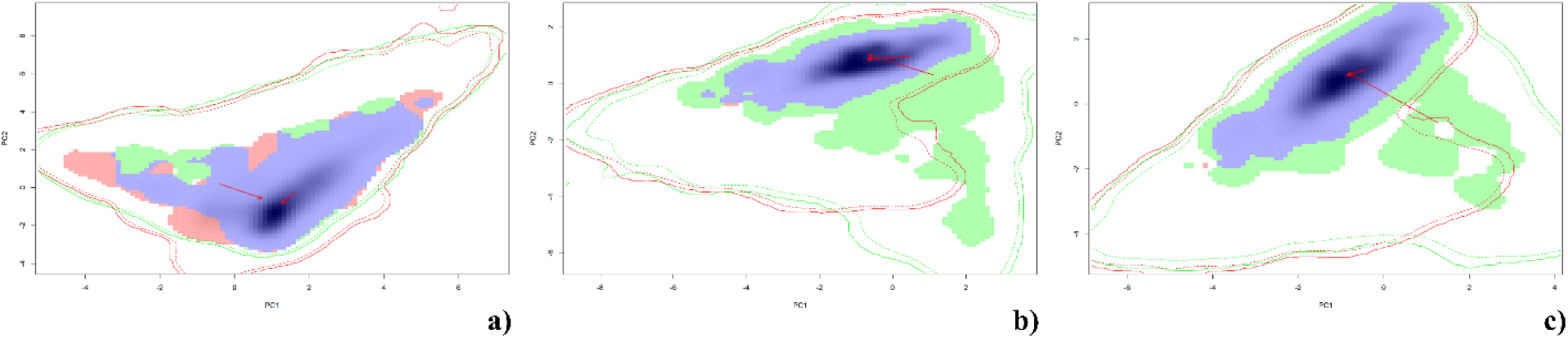
**a, b, c.** *Mimulus* niches in the European environmental space: a) *M. luteus* (green) and *M. guttatus* (red), b) *M. guttatus* and *M. robertsii* (red), c) *M. luteus* (green) and *M. robertsii* (red)

### Ecological niche of the parental taxa: relationship between invasive and native populations

#### *Mimulus guttatus* (common yellow monkeyflower)

Although our results indicate that the ecological niche of invasive populations of *M. guttatus* in Europe is similar to native populations, we found that there is an overall low niche overlap between them. The low overlap is associated with a large amount (61%) of niche unfilling, meaning that the invaded niche covers only a fraction of the environmental variability present in the native one (Supplementary Materials Figure 4a), which is consistent with niche conservatism for introduced populations of *M. guttatus* in Europe. Accordingly, previous studies on *Mimulus* species showed that *M.guttatus* native populations present a broad climatic niche (Ferris et al. 2014; Grossenbacher et al. 2014; Sheth and Angert 2014) and previous work on other systems has found that niche unfilling is more common than niche shift in terrestrial plants because the populations in the new environment occupy only a subset of the native environmental range (Petipierre et al. 2012; Strubbe et al. 2013; Guisan et al. 2014). Consistent with the idea that invasive European populations do not presently occupy the full range of environments covered in the native range, the projection of the native population niche into Europe shows highly suitable niche areas outside the current invasive distribution (Fig. 2a), whereas the species occurs mainly in the norh-western Europe and the British Isles.

The re-projection of the European invasive niche into the native areas highlight only a portion of the north-west part of the American continent as suitable for the species, and in particular the Aleutian Islands. Recent genetic analyses of UK populations of *M. guttatus* suggested a geographic origin population in the north Pacific (Puzey and Vallejo-Marín 2014; Pantoja et al. 2018). The niche analyses findings are also consistent with historical records, which indicate that one of the first *M. guttatus* specimens recorded in the UK originated from material collected in the Aleutian Islands in Alaska (Sims 1812; Pennell 1935, p. 116). As shown in Supplementary Materials Figure 3, the PCA made on the occurrences’ climatic data for three *M. guttatus* population (UK, occurrences further north than Haida Gwaii Islands and occurrences further south), showed the UK population closely correlated to the northern north American population and having only a restricted range of climatic conditions. Our findings support the niche conservation of *M. guttatus* in the European invaded areas and are also in accordance with the genetic analysis that suggest the North Pacific as the origin source of the European populations. To our knowledge, the use of ecological niche modelling to predict the geographic origin of invasive populations assuming the conservation of the realized niche and using records from the invasive range was applied in very few instances (e.g. Hardion et al. 2014). Hardion et al. (2014) used the distribution of invasive populations of *Arundo donax* (giant cane) in the Mediterranean region to infer the source of origin of this global invasive to an area in the Middle East, refining the hypothesised source of origin to southern Iran and the Indus Valley.

The ecological niche of the invasive New Zealand populations seemed not equivalent and similar by chance when compared to the native niche and the European one. These findings, coupled to a low D scores and niche dynamics suggesting niche unfilling (61%) in the fist case and both niche unfilling (24%) and expansion (48%) in the second one, seemed to indicate a niche shift in the ecological niche of the invaded population (Supplementary Materials Figures 4b, 5). The difference in ecological niche detected between European and New Zealand populations could reflect origins from source populations with slightly different climatic characteristics and/or be caused by post-colonisation changes allowing fine-tuning niche evolution. The timing of the naturalisation of *M. guttatus* in New Zealand (1878; Owen 1996) is compatible with a colonization event from British plants, which had become widespread in the UK by the mid 1800’s. Alternatively, New Zealand could have been independently colonised directly from the native range or from other populations, perhaps as part of the horticultural trade or seed exchange between botanic gardens. These inferences should be carefully interpreted considering 1) New Zealand’s population small size (only 19 occurrences), 2) that both niche dynamics analysis reported niche unfilling, and 3) that the PCA made on the environmental predictors highlighted non-analog conditions in the invaded range. However, ongoing genetic work indicates that at least some New Zealand populations can be traced back to the UK (Vallejo-Marin et al. unpublished).

#### *Mimulus luteus* (blood-drop-emlet monkeyflower)

The ENM of non-native populations of *M. luteus* showed suitable areas mainly in the British Isles, which is consistent with the current distribution of this taxon. The invasive niche is similar but non-equivalent to the native one, with evidence of both niche unfilling (35%) and expansion (16%; Supplementary Materials Figure 6a). While these findings statistically reject the niche conservation hypothesis, it is important to consider, as in the case of *M. guttatus*, that observed differences found between the native and invasive niches could reflect subsampling of the variation in environmental niche among populations in the native range. In the native range, *M. luteus* presents different morphological varieties that are partly geographically structured, although it is unknown whether they occupy different ecological niches (Carvallo and Ginocchio 2004). To date there is no genetic evidence of the source of origin on non-native opopulations of *M. luteus.* Based purely on niche similarity, we would predict that the source of the invasive European populations of *M. luteus*, if there is a single one, might came from the predicted highly suitable area in northern Patagonia.

#### Comparison between *M. guttatus* and *M. luteus*

Interestingly, the comparison between the niche of the parental taxa in both their native and European ranges, showed niche equivalency between the two species and niches more similar than expected by chance. The two species seems to experience similar environmental requirements in both ranges, although the niche overlap between *M. guttatus* and *M. luteus* is lower in the American range than in the European one (D = 0.384 and D = 0.734, respectively). In fact, as shown in Supplementary Materials Figure 6b and by the unfilling expansion index (34%), the two niches of these two taxa do not fully overlap in their native ranges, as expected for different species and given the different sample size between the two taxa. Closely related species often show similar but not equivalent niches (e.g. Aguirre-Gutiérrez et al. 2015; Dagnino et al. 2017), but our findings suggest that these two species have colonized similar habitats in the invaded range.

## Conclusions

This study provided the first ENM and niches comparisons of these three closely related monkeyflower taxa in their native American and invasive ranges. Niche conservation was supported for European *M. guttatus* population as well as for the comparison between the *M. luteus* (Inv.) population and the hybrid. On the contrary, the evidence of niche shift found must be interpreted with caution due to a) non analog climatic conditions between ranges (Guisan et al. 2012); c) niche unfilling dynamics and c) the small size of both invasive and native populations (for New Zealand s *M. guttatus* and *M. luteus* (Nat.), respectively). Retrospective ecological niche modelling allowed to predict the geographic origin of *M. guttatus* (Inv.) European populations, highlightig Aleutian Islands as the potential source of the first *M. guttatus* specimen recorded in Europe, as also supported by recent genetic analysis. However, retrospective ENM effectiveness strongly depend on the equivalency of both niches and the presence of analog environmental condition in both ranges. *M. robertsii*’s ecological niche showed a high degree of overlap to both progenitors in the environmental space, although it appears more related to *M. luteus* rather than *M. guttatus*. The coexistence of the species occupying similar environmental niches in their invasive range might be possible due to biotic factors not included here such as dispersal strategies that might be affected by future climate scenarios, increasing the species distribution. Future development of ecological niche models that include a mechanistic approach for the species should be considered in order to study more accurately the niche differentiations of the species by hybridization and invasion.

## Acknowledgements

We thank the Botanical Society of the Britain and Ireland (BSBI) for providing access to their monkeyflower records. DDR was supported by an ERASMUS+ 2016-2017 grant provided by the European Commission. MVM was supported by the University of Stirling.

## Supplementary Materials

**SM1.**
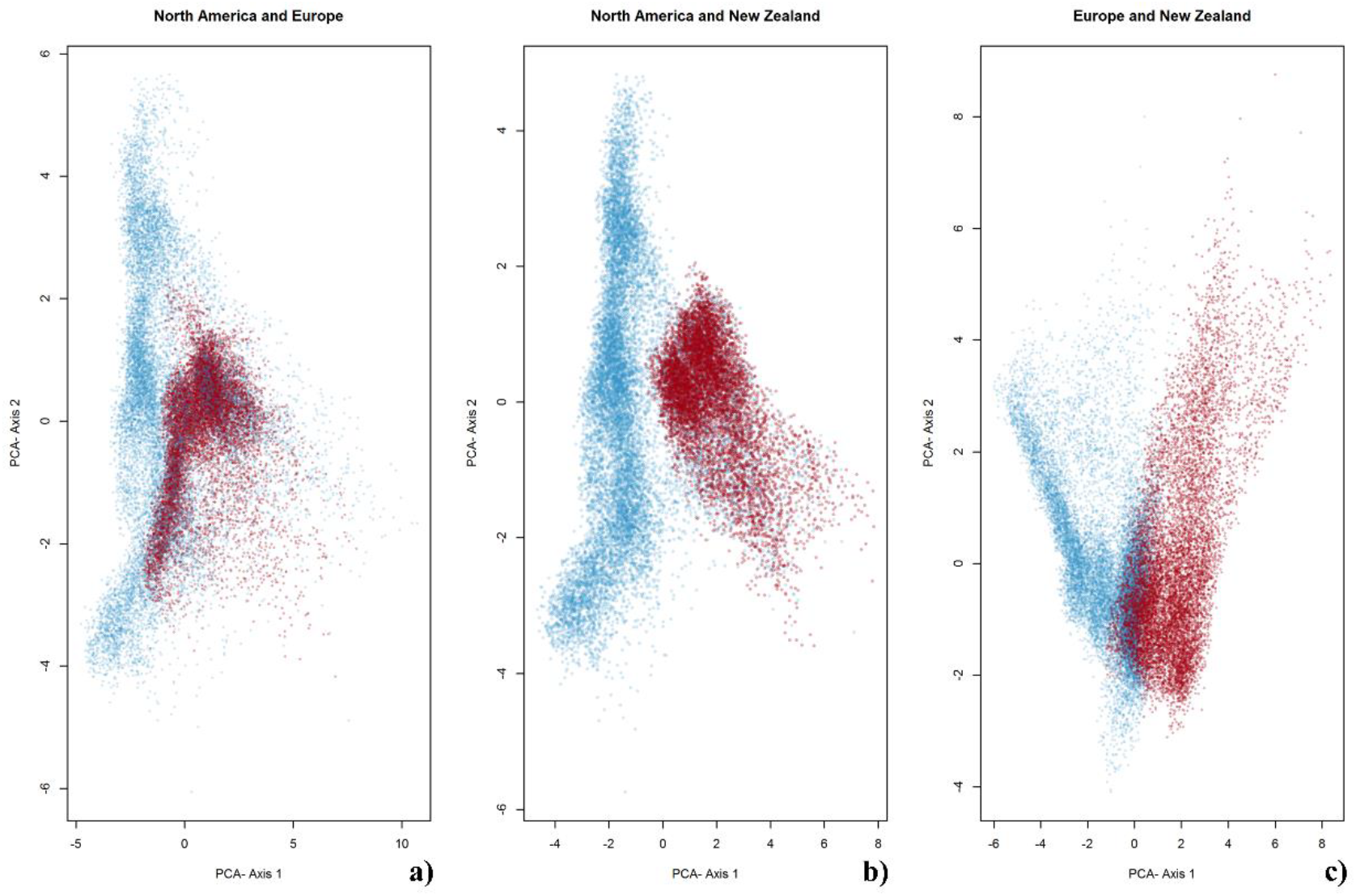
PCA on the climatic predictors for *M. guttatus* in a) Blue = North America, Red = Europe; b) Blue = North America, Red = New Zealand; c) Blue = Europe, Red = New Zealand.

**SM2.**
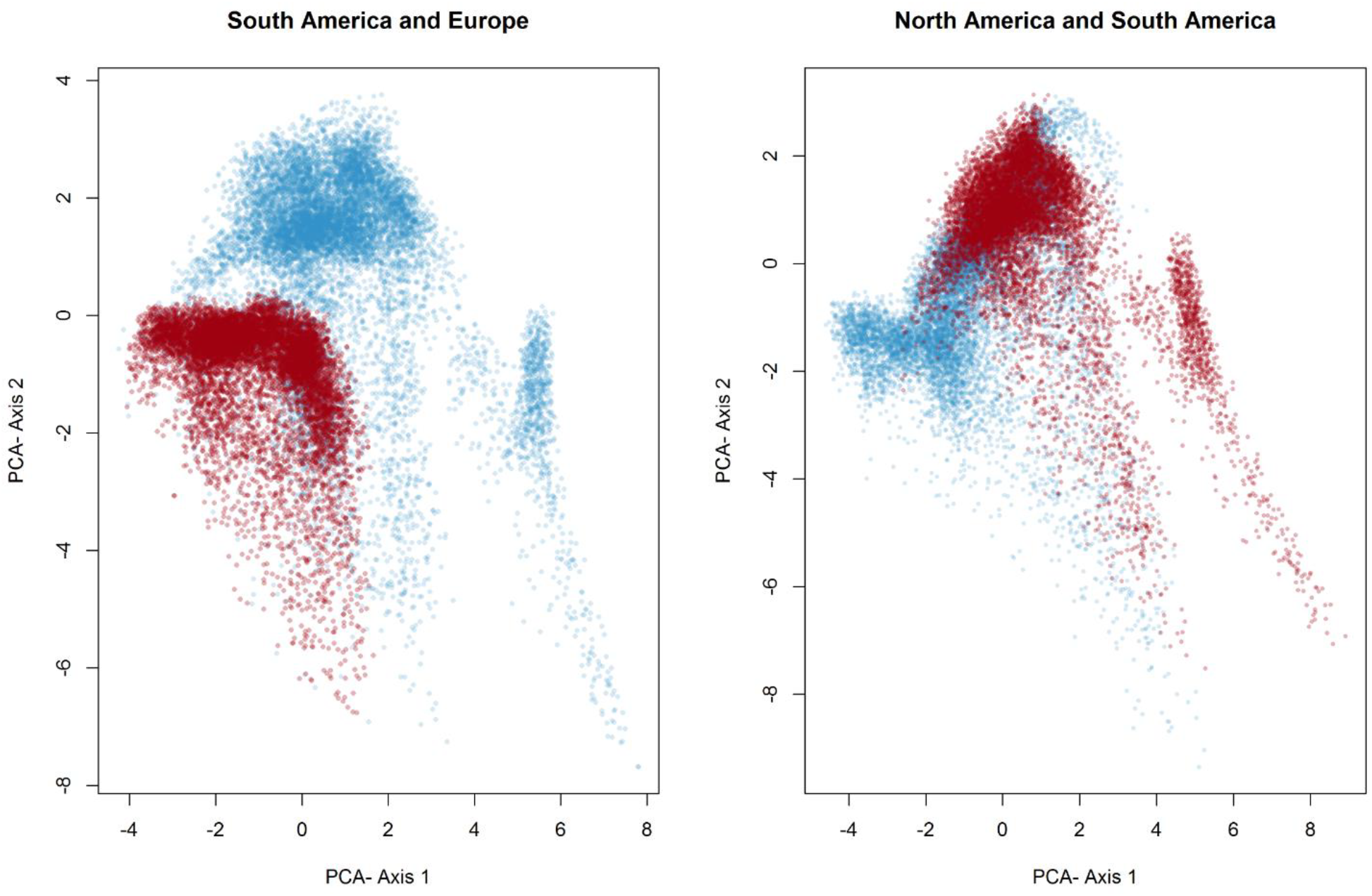
PCA on the climatic predictors for a) *M. lutes* in Blue = South America, Red = Europe; b) Blue = *M. guttatus* in North America, Red = *M. luteus* in South America.

**SM3.**
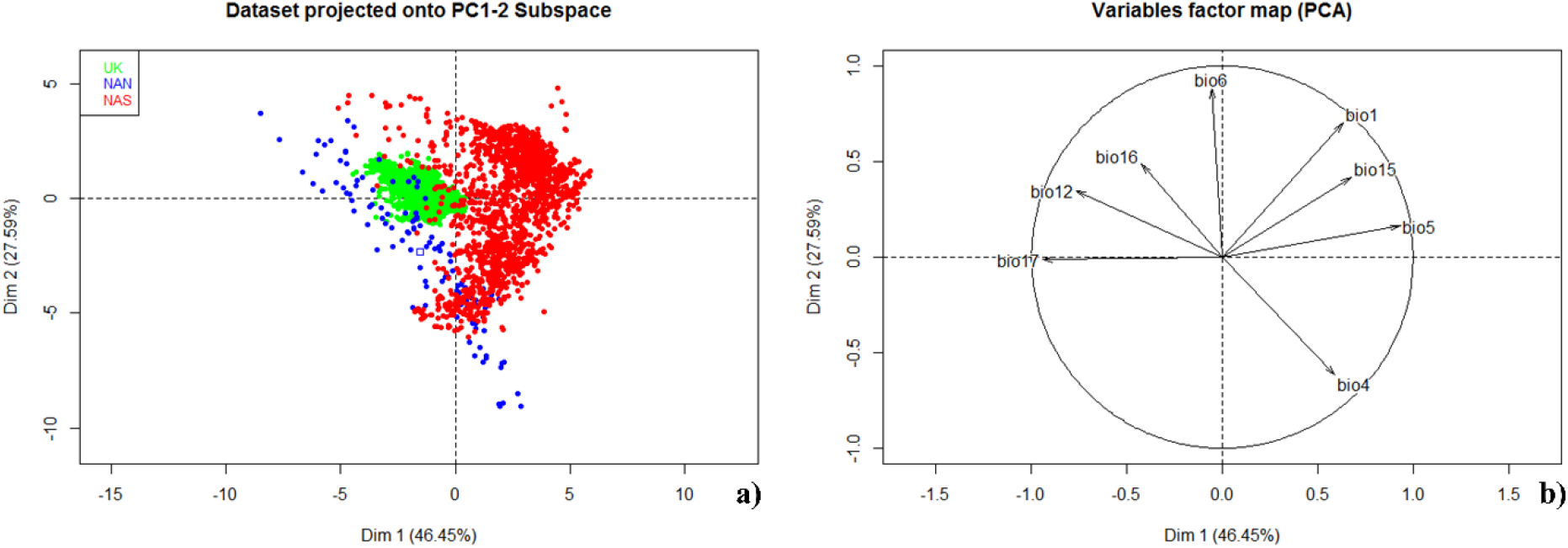
the PCA made on the climatic data for three *M. guttatus* population (UK, occurrences further north than Queen Charlot Island and occurrences further south than Queen Charlot Island). a) Individulals plot, b) variables plot. UK: M. guttatus occurrences in UK; NAN: M. guttatus occurrences further north than Queen Charlot Island; NAS: M. guttatus occurrences further south than Queen Charlot Island

**SM4.**
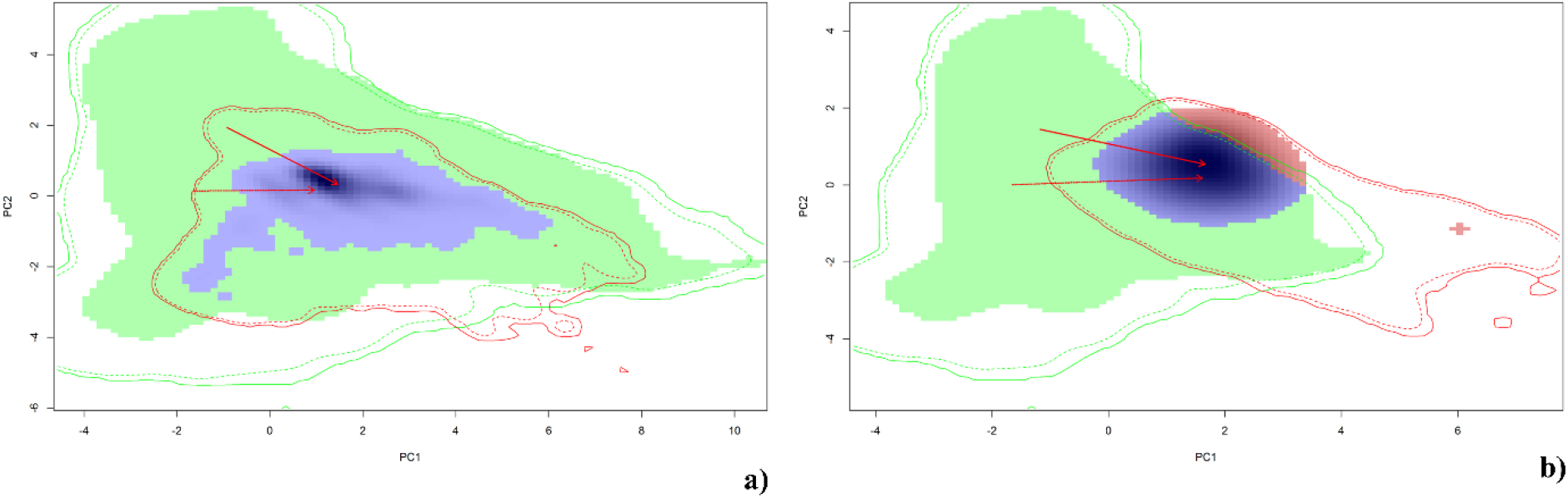
*M. guttatus* niches in the environmental space: a) Native niche (green) and Invasive European niche (red), b) Native niche (green) and Invasive New Zealand niche (red)

**SM5.**
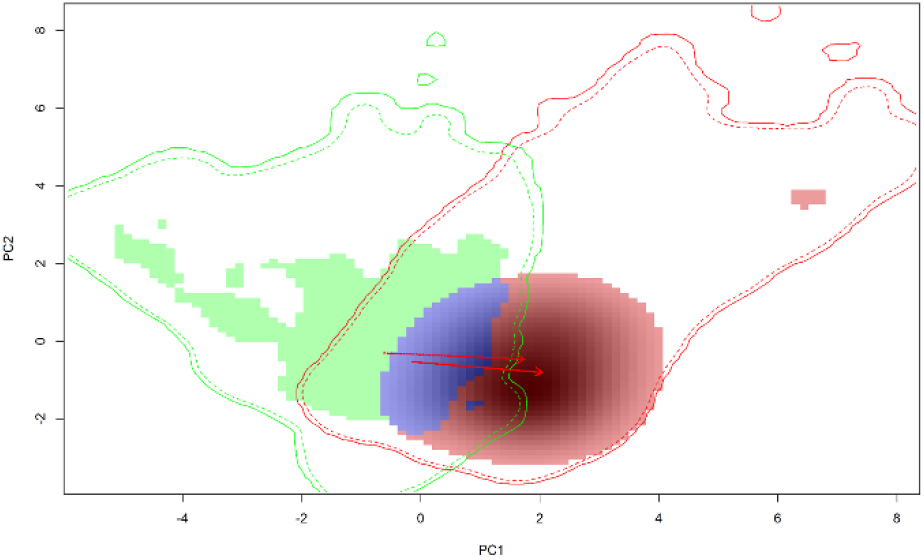
*M. guttatus* invasive niches in the environmental space: European (green) and New Zealand niche (red).

**SM6.**
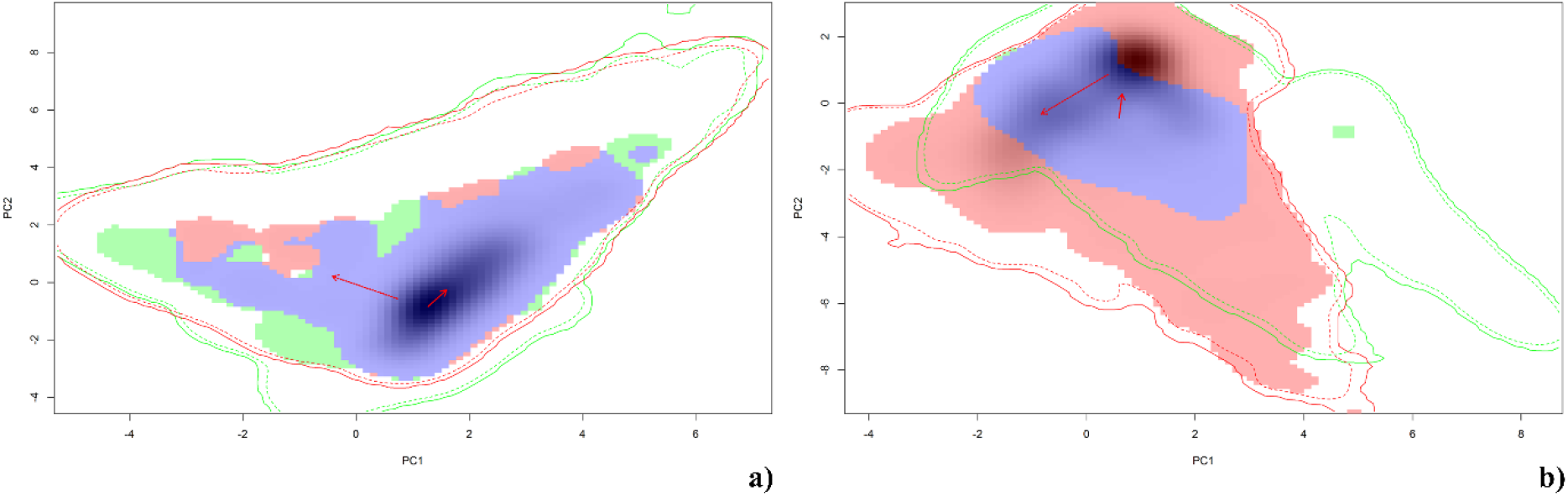
*M. luteus* niches in the environmental space: a) Native niche (green) and Invasive European niche (red), b) Native niche (green) and *M. guttatus* native niche (red)

